# The kinematics of 3D arm movements in sub-acute stroke: impaired inter-joint coordination caused by both weakness and flexor synergy intrusion

**DOI:** 10.1101/2022.12.03.518692

**Authors:** Inbar Avni, Ahmet Arac, Reut Binyamin-Netser, Shilo Kramer, John W. Krakauer, Lior Shmuelof

## Abstract

It has long been of interest to characterize the components that generate the motor control abnormalities in the arm after stroke. One approach has been to decompose the hemiparesis phenotype into negative signs, such as weakness, and positive signs, such as intrusion of synergies. Here, we sought to identify the contributions of weakness and the flexor synergy to motor function and impairment, as defined by kinematic and clinical scales, respectively, in sub-acute stroke using two 3D arm tasks that differed in their requirement for elbow extension. Thirty-three sub-acute post-stroke participants and sixteen healthy controls performed a cup-to-mouth task, requiring shoulder and elbow flexion (within flexor synergy), and a reaching task, requiring shoulder flexion and elbow extension (out of flexor synergy). Using markerless 3D pose-estimation, we analyzed upper limb kinematics to assess overall task performance and intrusion of pathological synergies. Weakness was measured using a grip dynamometer. Performance in both tasks was impaired to a similar degree in the stroke participants compared to controls. Subsequent analysis of coordination patterns between the elbow and the shoulder joints revealed intrusion of synergies in the reaching task based on the time spent within a flexion-flexion pattern (flexor synergy proportion) and the correlation between shoulder and elbow angles when the shoulder was flexing (flexion synergy strength). Regression analysis indicated that the significant predictors of poor task performance were weakness and flexor synergy intrusion. Notably, the Fugl-Meyer Assessment was abnormal even when just weakness caused the impairment, which means that caution is required when using this scale to quantify synergies. We conclude that both weakness and synergy intrusion contribute to impaired coordination of the elbow and shoulder joints in the sub-acute post-stroke period. This study shows that careful kinematic analysis of naturalistic movements is required to better characterize the components of upper limb impairment after stroke.

## Introduction

Approximately 80% of stroke survivors experience motor deficits, typically in the form of hemiparesis.^1,2^ Between 50% to 60% of patients with an initial arm paresis have arm disability at six months^3^ and one year.^4^ We have argued that the examination of the process of true motor recovery requires a focus on the impairment level.^1^ Motor impairment in the arm after stroke has multiple components: weakness, reduced motor control or dexterity, sensory loss, spasticity, and intrusion of pathological synergies.^1,5,6^ These components may be associated with distinct neural substrates and recovery profiles.^1,7,8^ Twitchell and Brunnstrom first described pathological flexor and extensor synergies in the context of patterns of limb movement at the joint level that emerge after stroke, which are comprised of either obligatory flexion or extension.^5,6^ In the clinical setting, a widely used motor impairment measure is the Fugl-Meyer Assessment (FMA), which quantifies the abilities of the participants to make isolated and coordinated joint movements on an ordinal scale.^9^ The FMA was designed to emphasize the contribution of pathological synergies, flexor and extensor, to post-stroke limb deficits. For example, in one maneuver, participants are required to flex the shoulder (0°-90°) while maintaining a straight elbow (0°). In this case, any flexion at the elbow would indicate the intrusion of a flexor synergy and lead to a lower score.

EMG can also be used to measure abnormal flexor and extensor muscle co-activation patterns during 2D isometric movements of the upper limb post-stroke.^10–12^ However, kinematic analysis is still required to quantify the effect of synergies as Brunnstrom (1966) and Twitchell (1951) described them, i.e., demonstrating their intrusion into natural movements. This is because tasks are kinematically defined: Did the participant make a normal-looking prehension movement or not? Indeed, kinematically normal movements can be made in the presence of significant co-contraction and high levels of stiffness.^13,14^ Thus, abnormal EMG is almost certainly necessary, but not sufficient, to generate abnormal joint kinematics; these need to be shown directly. In recognition of these issues, studies have designed reaching tasks that promote the emergence of synergies and then performed kinematic analyses. In one approach, participants are required to perform movements while bearing different shoulder loads,^15–17^ and in another, the kinematics of movements performed in and out of synergy are compared.^18^ For example, Zackowksi and colleagues (2004)^18^ showed that in chronic stroke, reaching kinematics in a task within flexor synergy were less impaired compared to a task that required elbow extension (out of flexor synergy). Notably, the stroke participants were also impaired in isolating movements around the wrist, elbow, and shoulder joints; flexion spilled over to the other joints, which led the authors to conclude that abnormal reaching kinematics in chronic stroke are driven by intrusion of the flexor synergy. While joint individuation has also been examined in sub-acute stroke,^19^ intrusion of pathological synergies during reaching has not. Here, instead of using the indirect measure of joint individuation, we devised two measures that directly detect flexor-synergy intrusion at the level of joint kinematics during 3D functional arm movements.

## Materials and methods

### Participants

Participants with either an ischemic or a hemorrhagic stroke (confirmed by imaging) were recruited by the Negev lab (a collaborative initiative of Ben-Gurion University and Adi Negev Nahalat Eran in Israel) between 2019 and 2022. Research protocols for both stroke and healthy participants were approved by Sheba Hospital Helsinki Committee and Ben-Gurion University Human Participants Research Committee, respectively. Only participants who were able to give informed consent were recruited. Additional inclusion criteria were: 1) Intact cognitive and motor control abilities before the incidence and 2) Sufficient active movement of the arm. Participants were excluded if they had a history of physical or neurological conditions that interfered with either the study procedures or the assessment of motor function (e.g., severe arthritis, severe neuropathy, Parkinson’s disease).

We analyzed data from 33 stroke participants in the sub-acute stage (1-8 weeks post-stroke, aged 65.1 ± 11.8) (Table 1) and 16 healthy controls (aged 68.8 ± 3.5). All participants were recorded performing the two tasks. Four recordings of two stroke participants performing the cup-to-mouth task with their right arm, and two others performing the same task using their left arm were missing due to technical difficulties. The motor component of the upper limb FMA was measured in all participants, as well as impairment measures of spasticity (Modified Ashworth Scale) and weakness (grip dynamometer).

**Table 1.**
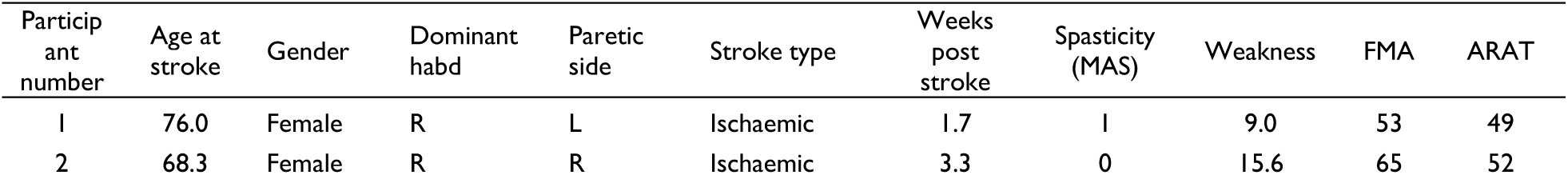
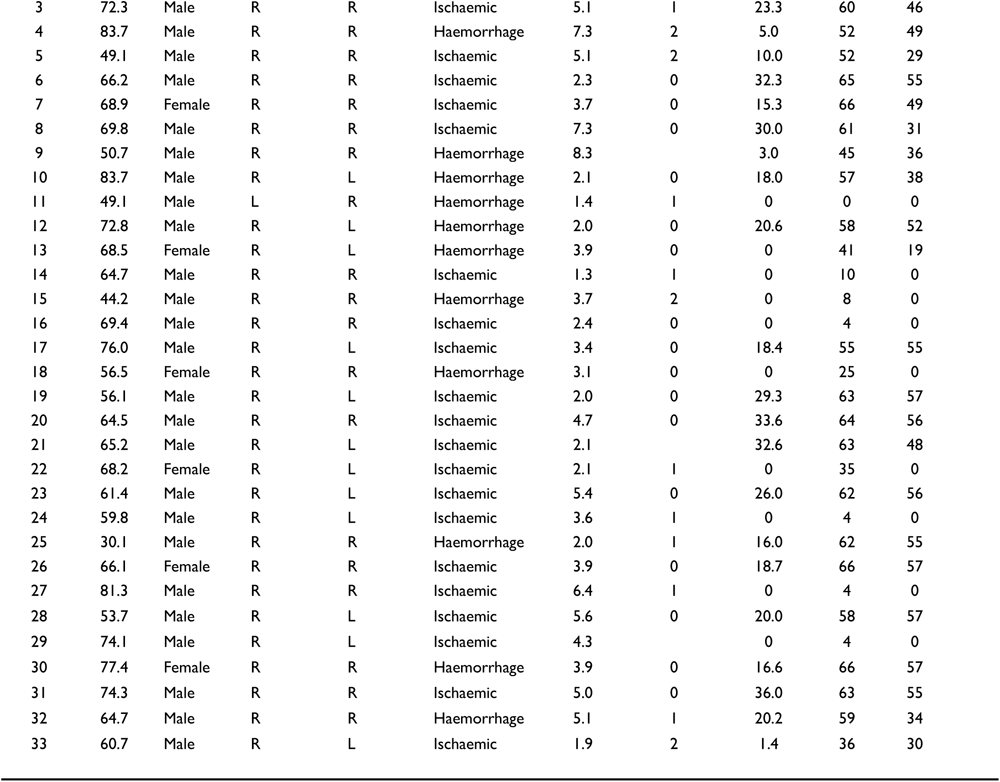
Characteristics of participants in the sub-acute phase after stroke.

### Experimental design

Two 3D arm tasks were recorded using markerless kinematics,^20^ a cup-to-mouth task that required shoulder and elbow flexion, and a reaching task that required shoulder flexion and elbow extension. In the cup-to-mouth task, participants were instructed to perform a simulated cup-to-mouth motion from a side table to their mouth, holding a plastic cup in their hand (taken from an ARAT kit). In the reaching task, participants were instructed to perform upward and forward reaching movements towards a suspended target. The instruction was to complete ten trials of each task. To compare the two tasks, in the cup-to-mouth task, only the movement segments of reaching from table to mouth were examined.

To quantify movement, we used a novel approach for analyzing markerless 3D kinematics (DeepBehavior)^20^ using a convolutional neural network algorithm (OpenPose)^21^ that was trained to detect 57 key points in the human body in each video frame (Fig. 1).

**Figure 1.**
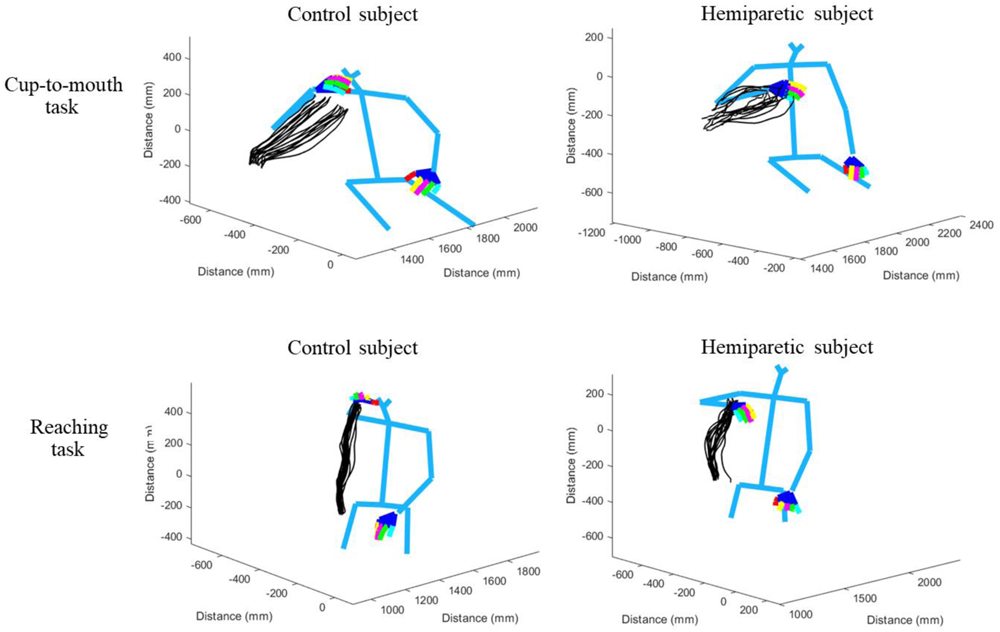
Kinematics of the human arm and body during two 3D tasks. Estimation of 3D movements in two participants: blue lines connect the model’s joint positions, and the movement trajectories are marked by black lines (each participant performed ten repetitions of the task). Trajectories are shown for a healthy control participant (left column) and a stroke participant (right column) performing the cup-to-mouth task (top row) and the reaching task (bottom row).

### Recording setup

The tasks were recorded using a custom-made system comprising two high-speed cameras (150 frames per second, 1280x1024 pixels, Blackfly S Color 1.3 MP USB3 camera with a Fujinon 1.5MP 6mm C Mount lens), set on a custom-designed aluminum camera holder with a 66° angle between their axes. In this setup, cameras were positioned 120 cm in front of the participant, at a height of 95 cm, and placed at a set 45° angle towards the participant, which allowed us to produce the 3D kinematic data of 57 body key points. In the cup-to-mouth task, participants faced the cameras at a 45° angle with the moving arm closer to the camera lenses and a side table with an empty cup placed to their side. In the reaching task, participants were recorded from the frontal angle while facing the cameras and reaching up towards a suspended object (∼1.5 m above ground). Each participant attempted to perform ten iterations of each task in each hand (in separate blocks). Stroke participants that could only partially execute these tasks were included in the analyses if minimal paretic arm movement was detected at least twice in each task. All participants in our study were able to perform shoulder elevation (as indicated by a clinician). During the recordings, no markers were placed on the participants as the analysis algorithm enables markerless detection of joint positions.

### Impairment and functional measures

FMA,^9^ Action Research Arm Test (ARAT),^22^ spasticity (Modified Ashworth Scale, MAS),^23^ and weakness (grip dynamometer) scores were collected from all participants. Neither MAS nor weakness scores were collected when the arm was flaccid, or the participant couldn’t perform power grip. The values obtained using the grip dynamometer were normalized into z-scores according to the age, gender, and dominant hand of each of the participants.^24^

### Data analysis

The recordings resulted in two synchronized videos from two cameras. Each video was passed through the OpenPose algorithm to detect joint positions. Then, the corresponding 2D positions of joints from each video were stereo-triangulated to obtain the estimated 3D position.^25^ To do this, a prior calibration using a checkerboard was obtained. This resulted in a list of 3D positions of all joints. These data were smoothened using a Savitzky-Golay filter with a window size of 57 and a polynomial degree of 3. Then, the joint tangential velocities were calculated. Movements were segmented based on the wrist velocity profiles (movement start and end were defined based on the crossing point of 10% of the peak velocity). Peak velocity detection and segmentation were automatic but verified and adjusted manually. The main performance measures were extent, peak velocity, movement duration, and smoothness. The extent was defined as the radial 3D position at the end of the reaching/cup-to-mouth task compared to the start position, movement duration, and peak velocity. Smoothness was defined as the minus log of the normalized integrated squared jerk (see equation 1).

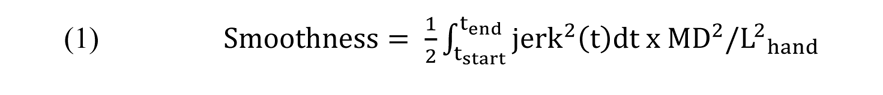

Joint angle data was calculated using an intrinsic (anatomical) coordinate system (angle calculated in relation to a specific joint; shoulder flexion angle is defined as the angle in degrees between the ipsilateral elbow joint, ipsilateral shoulder joint, and contralateral shoulder, projected onto the sagittal plane – defined by the torso and shoulder vectors created by the 3D model of the participant; elbow extension angle is defined as the angle in degrees between the ipsilateral wrist, elbow, and shoulder joints).

Flexion-flexion coordination was quantified based on the angular velocity of the elbow and shoulder joints in two ways: 1) flexion-flexion proportion was calculated as the time spent while simultaneously flexing the elbow and shoulder, divided by the total time of the movement, and 2) flexion-flexion strength was the Pearson’s correlation coefficient of the shoulder flexion and elbow extension angles, during the largest segment in the movement that the shoulder was flexing (determined based on their angular velocities). To deal with the skewed distribution of this measure, we performed a fisher transformation by calculating the inverse hyperbolic tangent (arctanh) of the correlation coefficients. Data analysis was performed using a custom-written code in MATLAB.^26^

### Statistical analysis

Between-group differences (control, non-paretic stroke, and paretic stroke) were assessed using two-sample, two-tailed t-tests with unequal variance. Cohen’s d was used to determine the effect sizes of differences between groups, and the Bayes Factor was calculated to describe the strength of evidence for the alternative hypothesis vs. the null hypothesis. Additionally, two-way ANOVA analyses were performed to assess the differences in kinematic measures between groups (control and paretic stroke) and across tasks.

Furthermore, the contribution of different impairment measures (spasticity, weakness, and intrusion of synergy) to performance measures (e.g., smoothness and extent) in the data of stroke participants was assessed using linear regression analysis. Statistical analysis was performed via MATLAB^26^ and JASP.^27^

### Power analysis

Based on the final sample containing 33 stroke participants and 16 age-matched controls, the power to identify significant differences across groups (three groups: controls, non-paretic arm measures, and paretic arm measures of the stroke group), in a two-way ANOVA assuming a medium effect size of f=0.4 across groups, is 89%. Furthermore, our correlation analysis has a power of 94% to identify an effect of ρ=0.5. The multiple linear regression analysis has a power of 92% to determine an effect of f^2^ = 0.3 with three predictors in the model. Statistical power was computed using G*power version 3.1.9.4.^28^

## Results

### Performance was abnormal in both 3D arm tasks

To test for a possible effect of a flexor synergy on arm movements after stroke, we had participants perform a reaching task that required flexion at the shoulder and extension at the elbow (movement outside of flexor synergy) and a cup-to-mouth task that required flexion at the shoulder and elbow (movement within flexor synergy) (Fig. 1 and Supp Fig. 1-6).

Movements of stroke participants appeared slower and shorter, showed increased jerkiness, and sometimes involved compensatory strategies. To go beyond observation, performance on the two tasks was quantified by measuring each participant’s peak velocity, movement duration, movement extent, and smoothness.

Inferior performance in stroke participants was seen when comparing the paretic side with the performance of control participants in all measures and tasks (Fig. 2). When comparing the performance of the non-paretic side with the performance of age-matched control participants, no significant differences were observed in any of the measures of the cup-to-mouth task (*P* > 0.05). In the reaching task, the same comparison yielded significant differences in movement duration, peak velocity, extent and smoothness (P < 0.05) (Fig. 2 & Table 2). The stroke participants were also significantly weaker than the control group (t(32) = -9.47, *P* = 0, d = -36.2, BF10 > 10^6^).

**Figure 2.**
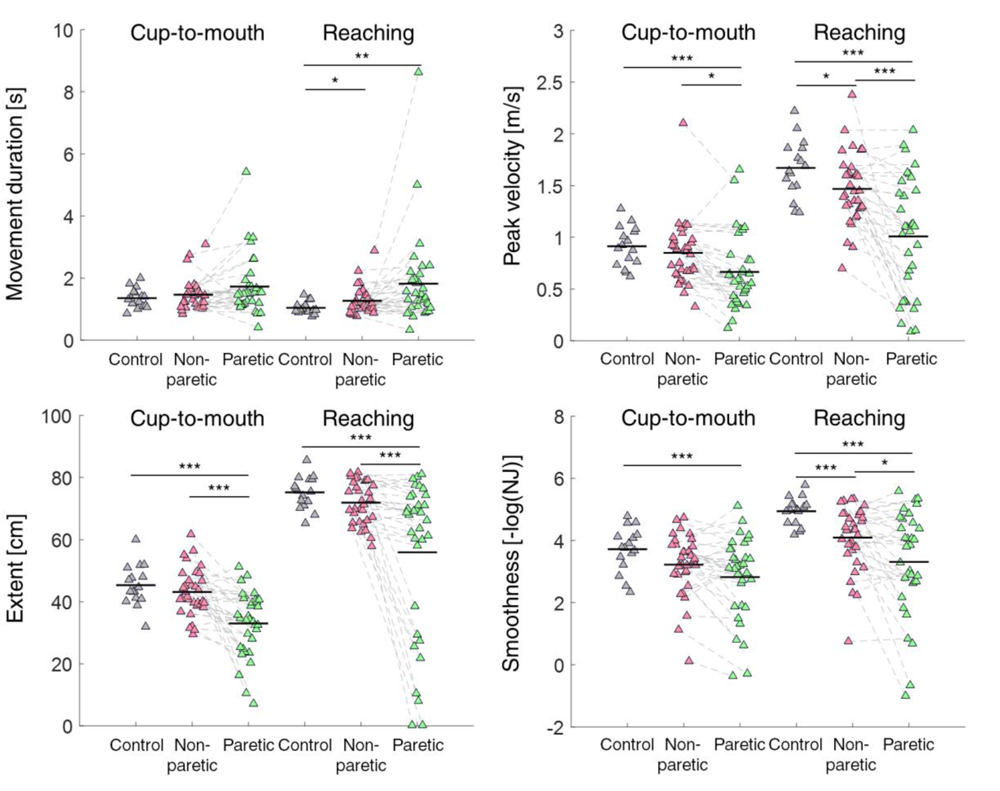
Kinematic measures of performance showed impaired performance in both the reaching and cup-to-mouth tasks after stroke. (**A**) Scatter plots of movement duration, (**B**) peak velocity, (**C**) extent, and (D) smoothness in each group, for both tasks. Each triangle represents the values of the measure for a single participant across all individual movements in the task. Horizontal lines represent averages. Significant differences across groups are denoted by asterisks (* = *P* < 0.05, ** = *P* < 0.01, *** = *P* < 0.001, two-sample t-test with unequal variance).

**Table 2.**
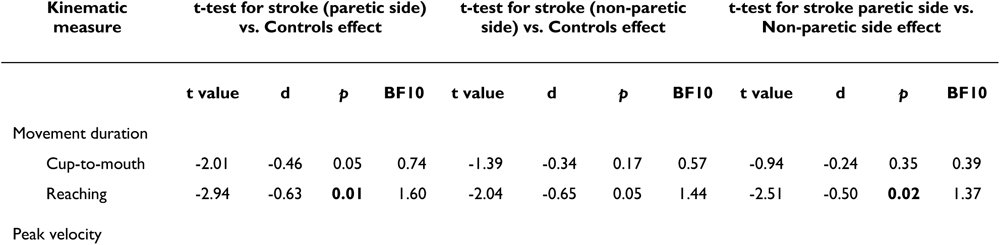
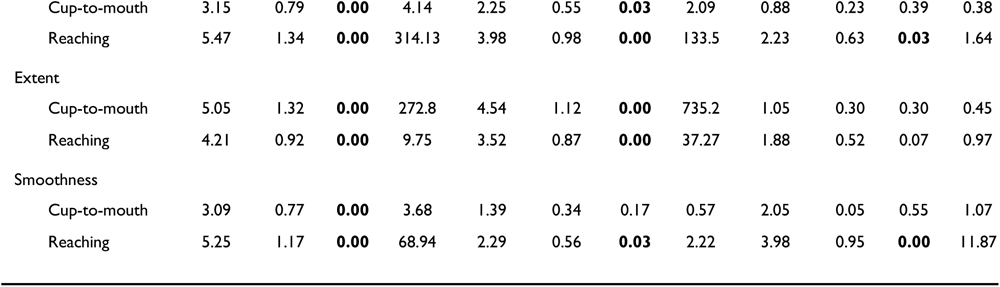
Statistical analysis of kinematic measures.

To test for synergy intrusion, we ran an interaction analysis between damage (paretic hand and control) and task (in and out of synergy). The interaction was significant for the measure of peak velocity (*P* = 0.02) (Table 3). Thus, participants were impaired compared to controls on both tasks, with a weak trend towards worse performance in the out-of-synergy task.

**Table 3.** Two-way ANOVA analysis of the interaction of group and task in kinematic measures.

| Kinematic measure | Sum of squares | df | F | p |
| --- | --- | --- | --- | --- |
| Peak velocity | 0.92 | 1,94 | 5.39 | **0.022** |
| Movement duration | 0.89 | 1,94 | 0.81 | 0.371 |
| Extent | 264.24 | 1,94 | 1.00 | 0.320 |
| Smoothness | 2.89 | 1,94 | 1.76 | 0.188 |

### Evidence for intrusion of the flexor synergy

To examine for intrusion of synergies into arm movements, we devised two novel approaches to measure the coordination of shoulder and elbow joint angles during performance. The first approach was to quantify the flexion-flexion pattern during arm movements in patients and healthy controls. To do this, we calculated the proportion of time that participants spent in flexion-flexion coordination during the movement by identifying the segments in which both the shoulder and elbow were flexing (“flexion-flexion proportion”) (see Fig. 3A). In neurotypical individuals, this proportion would reflect the amount of flexion-flexion that was required to complete the task. Any deviation from neurotypical behavior, whereby flexion-flexion proportion increased, was deemed a synergy.

**Figure 3.**
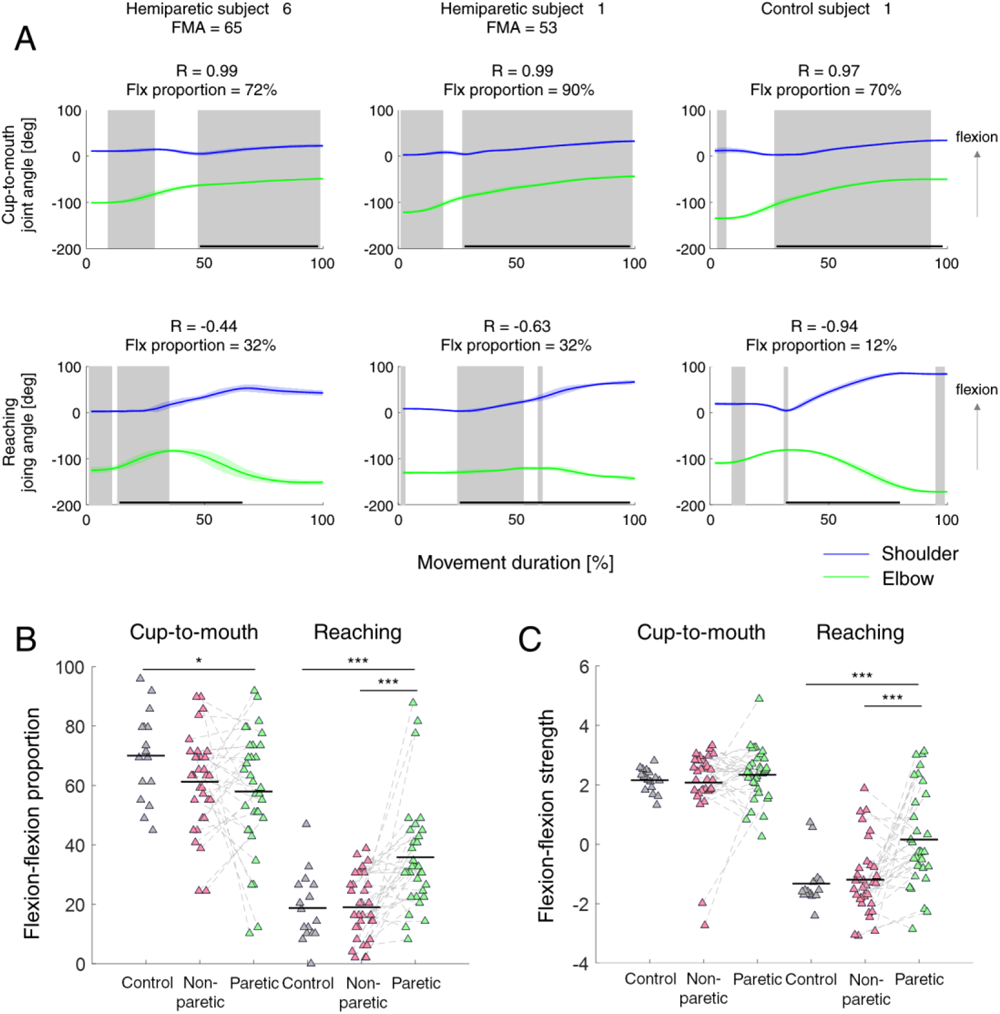
Two kinematic measures of flexor synergy intrusion. **(A)** Shoulder and elbow angular trajectories of two hemiparetic participants (left and middle) and a control participant (right) in both tasks: cup-to-mouth (top) and the reaching task (bottom). Gray areas represent the time points in which both joints were flexing, based on which the proportion of time spent in flexor synergy was calculated (flexion-flexion proportion). Horizontal black lines represent the longest segment identified of shoulder flexion. Based on this segment, Pearson’s correlation coefficient on the angular trajectories of the shoulder and elbow was calculated (flexion-flexion strength). Angular trajectories are averaged across all individual movements in the task. **(B)** Scatter plot of the percentage of time spent within flexion-flexion movement pattern in each group. **(C)** Scatter plot of the correlation coefficient of the flexion-flexion angles in each group. Correlation coefficients were measured only in segments of shoulder flexion. Each triangle represents the values of the measure for a single participant. Significant differences across groups are denoted by asterisks (* = *P* < 0.05, ** = *P* < 0.01, *** = *P* < 0.001, two-sample t-test with unequal variance).

Indeed, in the cup-to-mouth task, the control participants spent a greater percentage of time in flexion-flexion coordination (mean±std: 70.03 ± 15.1) than in the reaching task (mean±std: 18.75 ± 11.6). In addition, the proportion of time control participants spent in flexion-flexion coordination was highly affected by the task (*t* = 10.8, *P* = 0). Consistent with the intrusion hypothesis, this measure was greater for stroke participants in the reaching task (paretic vs. controls: t(43.4) = -3.97, *P* = 0, d = -1.04, BF10 = 24.17). The paretic side also showed increased flexor synergy compared to the non-paretic side in this task (t(32) = -4.27, *P* = 0, d = -1.12, BF10 = 163.6).

We next examined if stroke participants show a general increase in using a flexion-flexion pattern. Interestingly, this wasn’t the case - they spent significantly less proportional time than control participants in flexion-flexion coordination in the cup-to-mouth task (paretic vs. controls: t(38.66) = 2.34, *P* = 0.02, d = 0.65, BF10 = 1.75) (Fig. 3B). This observation suggests that intrusion of synergies interfere with the task only in those that require movements outside of synergy. The reduced flexion-flexion pattern in the cup-to-mouth task may be driven by other deficits.

Another way of assessing the level of intrusion of the flexor synergy was to assess the strength of dependency between the shoulder flexion and the elbow’s involuntary flexing. To do that, we analyzed the motion of the elbow when the shoulder was flexing. If abnormal synergy patterns interfered with the movement, this would mainly occur in the parts of the movement where the shoulder was actively flexing, enslaving the elbow into flexion. We, therefore, identified in each task the longest segment of shoulder flexion and calculated the Pearson’s correlation coefficient on the angular trajectories of the shoulder and elbow during that segment (Fig. 3C). This measure, termed “flexion-flexion strength,” would reflect the extent of shoulder flexion and elbow flexion dependency, and is expected to be positive in tasks requiring mostly a flexion-flexion pattern, and negative in tasks requiring a flexion-extension pattern. Here also, any deviation from neurotypical behavior, whereby the strength of the flexion-flexion pattern increased, was deemed a synergy.

Indeed, in the control participants, flexion-flexion strength was positive in the cup-to-mouth task (mean±std: 2.14 ± 0.40) and negative in the reaching task (mean±std: -1.34 ± 0.82). As with the flexion proportion measure, significant differences in flexion strength were observed between groups in the reaching task, with stroke participants exhibiting increased flexion-flexion strength (t(46.84) = -4.24, *P* = 0, d = -1.05, BF10 = 25.54), indicating flexor synergy intrusion. Consistent with the previous finding, flexion strength was not different between groups in the cup-to-mouth task, indicating that synergies intrude movements outside the flexor pattern (*P* > 0.23).

Notably, two control participants displayed slightly positive values in this measure in the reaching task (0.55 and 0.72). This can be explained by these participants’ idiosyncratic natural movement pattern that involved flexing their elbow before extending it to reach the target. Indeed, when calculating the Pearson Correlation coefficient of the entire movement, these participants produced negative values, indicating that overall, they extended their elbow while flexing the shoulder in the reaching task. Importantly, stroke participants showed increased flexor strength in their paretic arm compared to their non-paretic arm (t(32) = -3.79, *P* = 0, d = -0.94, BF10 = 48.51), indicating that high flexion strength reflects intrusion of synergy rather than an altered natural movement pattern.

To conclude, two direct kinematic measures of flexion-flexion coordination (proportion and strength) provide evidence of flexor synergy intrusion for movements requiring a flexion-extension coordination pattern.

### Inter-joint coordination was abnormal due to both weakness and synergy intrusion

So far, the results indicate impaired performance in both tasks, and intrusion of the flexor synergy in the reaching task. However, it is still unclear to what extent overall task performance can be attributed to the intrusion of a flexor synergy.

To test this, we sought to evaluate the differential contribution of different types of impairment to task performance. We applied regression analysis with kinematic measures as the dependent variables and the two synergy measures and grip dynamometry as independent variables (Table 4).

**Table 4.**
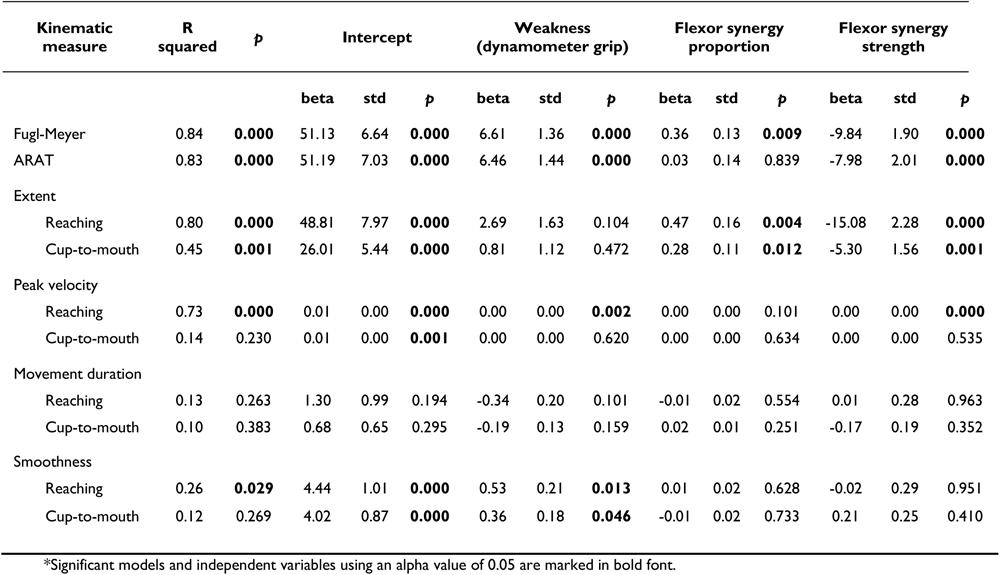
Regression analysis of different types of impairment measures.

In the reaching task, weakness was a significant predictor for the peak velocity and the smoothness measures, whereas flexor synergy predictors were significant predictors for extent (strenthgh & proportion) and peak velocity (strength) (Table 4). Notably, the flexor synergy intrusion measures were also significant predictors of extent in the cup-to-mouth task, but since the impairment in this case was associated with a reduction in flexion-flexion coordination, this predictor does not represent synergy intrusion.

Next, we evaluated the differential contribution of types of impairment to clinical measures by applying the same regression analysis approach with the FMA or ARAT as dependent variables and the two synergy measures (taken from the reaching task) and weakness as independent variables (Table 4). Both ARAT and FMA scores were significantly modeled by synergy measures and weakness (R^2^ = 0.83 and 0.84, respectively, *P* < 0.001) (Table 4).

To test whether spasticity plays a significant role in explaining these scores, we added a measure of spasticity (the Modified Ashworth Scale) to these models. Even though the Modified Ashworth Scale scores significantly correlated with these measures (FMA, ARAT, extent in both tasks and peak velocity in the reaching task) (Table 5), adding spasticity as an independent variable didn’t improve the fit of the regression models.

**Table 5.**
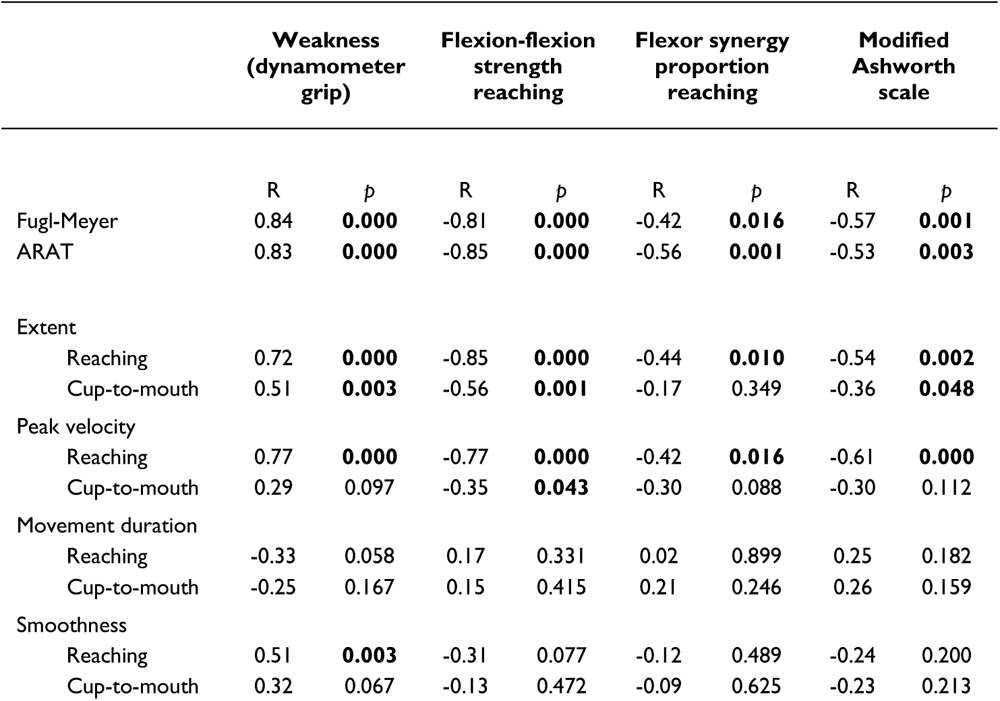
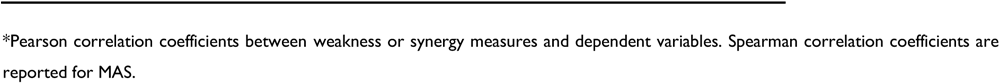
Correlation coefficients of different types of impairment and functional measures.

|  | Weakness (dynamometer grip) |  | Flexion-flexion strength reaching |  | Flexor synergy proportion reaching |  | Modified Ashworth scale |  |
| --- | --- | --- | --- | --- | --- | --- | --- | --- |
|  | R | p | R | p | R | p | R | p |
| Fugl-Meyer | 0.84 | **0.000** | -0.81 | **0.000** | -0.42 | **0.016** | -0.57 | **0.001** |
| ARAT | 0.83 | **0.000** | -0.85 | **0.000** | -0.56 | **0.001** | -0.53 | **0.003** |
| Extent |  |  |  |  |  |  |  |  |
| Reaching | 0.72 | **0.000** | -0.85 | **0.000** | -0.44 | **0.010** | -0.54 | **0.002** |
| Cup-to-mouth | 0.51 | **0.003** | -0.56 | **0.001** | -0.17 | 0.349 | -0.36 | **0.048** |
| Peak velocity |  |  |  |  |  |  |  |  |
| Reaching | 0.77 | **0.000** | -0.77 | **0.000** | -0.42 | **0.016** | -0.61 | **0.000** |
| Cup-to-mouth | 0.29 | 0.097 | -0.35 | **0.043** | -0.30 | 0.088 | -0.30 | 0.112 |
| Movement duration |  |  |  |  |  |  |  |  |
| Reaching | -0.33 | 0.058 | 0.17 | 0.331 | 0.02 | 0.899 | 0.25 | 0.182 |
| Cup-to-mouth | -0.25 | 0.167 | 0.15 | 0.415 | 0.21 | 0.246 | 0.26 | 0.159 |
| Smoothness |  |  |  |  |  |  |  |  |
| Reaching | 0.51 | **0.003** | -0.31 | 0.077 | -0.12 | 0.489 | -0.24 | 0.200 |
| Cup-to-mouth | 0.32 | 0.067 | -0.13 | 0.472 | -0.09 | 0.625 | -0.23 | 0.213 |

\*Pearson correlation coefficients between weakness or synergy measures and dependent variables. Spearman correlation coefficients are reported for MAS.

Thus, our regression results suggest that intrusion of synergies and weakness affect task performance and FMA and ARAT scores (Table 4).

It is important to note that the FMA clinical scores were related to synergy intrusion and weakness to a similar extent. It is perhaps not surprising that the FMA was also affected by weakness. Notably, when examining the participants with severe impairment as defined by the FMA (FMA below 45), we noticed that all 13 participants exhibited a significant level of weakness (lower than four standard deviations below the population mean). Most of these participants (11/13) also exhibited synergy intrusion (flexor synergy strength higher than zero) (Figure 4).

**Figure 4.**
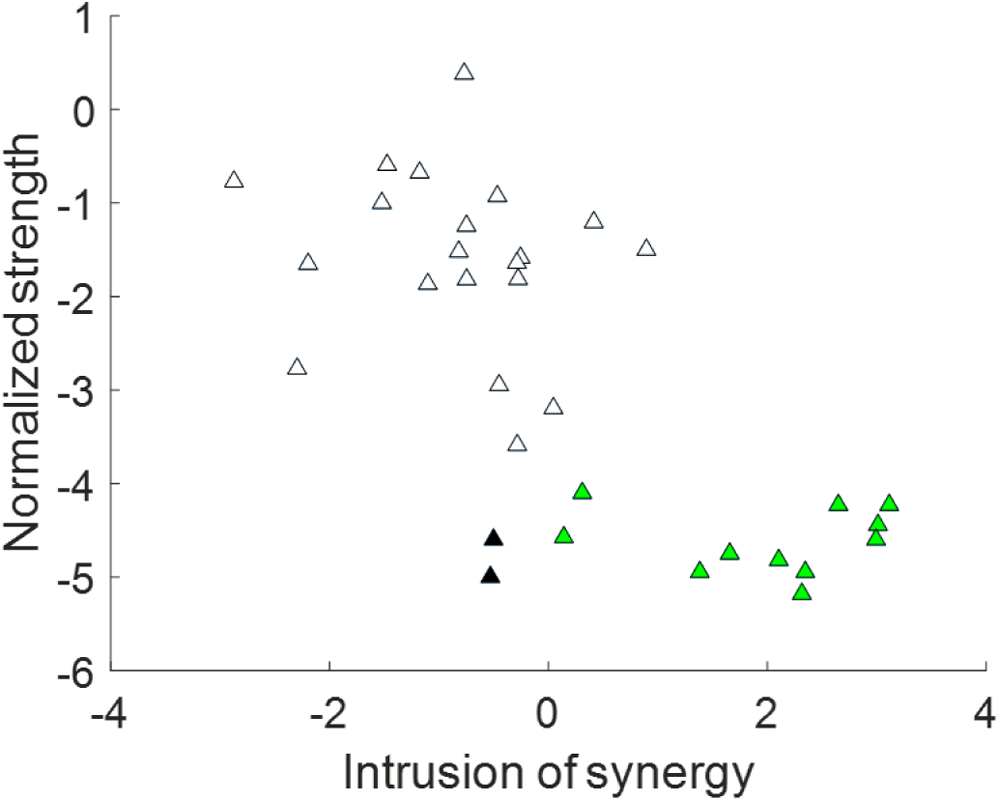
Impaired participants exhibit weakness and intrusion of synergies. Scatter plot of the weakness measure compared to the flexor synergy strength measure taken from the reaching task. Each triangle represents the values of the measure for a single participant across all individual movements in the task. Triangles of participants with lower FMA scores than 45 are colored: green for participants with intrusion of flexor synergy and weakness and black for participants with weakness but without intrusion of flexor synergy.

Looking from another angle, thirteen participants displayed positive values for flexion-flexion strength. Only two of these participants had normal strength and a higher score on the FMA (> 45) (Figure 4).

### Sources of motor impairment through observation

Though the contributions of flexor synergy and weakness were detected in the quantitative analysis, it is important to note that not all upper limb impairment can be attributed to pathological synergies. To characterize the unique impairment profiles of the stroke participants, a group of researchers and clinicians examined video clips for each participant with stroke.

The videos of participants were watched by a neurologist (JWK) and two occupational therapists (RBN, SD). One of the dominant problems noted by the clinicians was weakness-induced compensatory movements around the shoulder. Specifically, weakness was evident in the difficulty participants had anteriorly flexing the shoulder and extending the elbow to its full capacity (Fig. 5). Compensatory abduction and hiking of the shoulder led to internal rotation of the arm, with the result that the elbow fell into flexion (Fig. 5A). Weakness was also apparent in wrist drop (Fig. 5A). The clinicians also noted evidence for muscle adhesion and scapula impingement. Therefore, we argue that weakness and peripheral secondary impairments made independent contributions to inter-joint coordination deficits that went beyond synergy intrusion.

**Figure 5.**
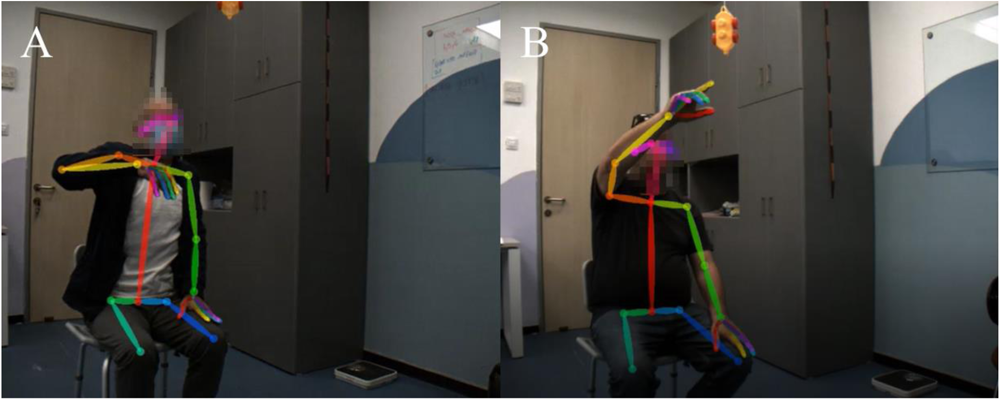
Examples of participants performing the reaching task. Two participants are presented at the end of the upward movement in the reaching task, demonstrating their difficulty completing it. Movies depicting the 3D model of the participants performing the task are also available in the supplementary material (Supp Mov. A and B, corresponding to each of the participants presented in this figure).

## Discussion

We sought to examine the underlying components of motor impairment after stroke with a novel approach for assessing the intrusion of the pathological flexor synergy to 3D arm movements using markerless pose-estimation. We found that intrusion of the flexor synergy and weakness both contributed to poor kinematic performance in two 3D functional arm tasks. Abnormal clinical measure scores in the sub-acute stage were also attributable to weakness and synergy intrusion.

Here, we demonstrated the benefits of using 3D kinematics, which allows a suitably granular analysis of reaching performance in naturalistic tasks. Performance was abnormal, with significant differences in all kinematic measures (Figure 1 & Table 2). Assessing kinematics in two tasks that differentially stress out-of-synergy movements allowed us to assess the presence of synergies, as has been done previously.^18,29^ Levin (1996) showed that inter-joint coordination deficits during 2D planar arm movements in participants with chronic stroke were invariant to reaching direction, suggesting that intrusion of synergies is not a major contributor to 2D post-stroke reaching deficits.^29^ In contrast, in chronic stroke, Zackowski and colleagues showed synergy intrusion by demonstrating greater deficits in 3D reaching requiring movement outside the flexor synergy than within the flexor synergy. They supported this conclusion by showing that participants with stroke could not make isolated movements of the wrist, elbow, or shoulder joints.^18^ Our kinematic analyses revealed slightly greater deficits in the out-of-synergy task and a mildly significant interaction between the group and task for peak velocity (Table 3), which are not consistent with the results of Levin et al.. Furthermore, our coordination measures of synergy provide additional support for the intrusion of synergies during the sub-acute phase after stroke.

A crucial benefit of kinematic analysis is that it allows the quantification in functional tasks of both negative symptoms, such as loss of dexterity and the effect of weakness, and positive symptoms, such as intrusion of synergies. ^29–31^ We developed two measures for flexor synergy intrusion during upper limb movements: “proportion” and “strength.” The proportion measure sought to capture the effect of synergies on the overall coordination of the movement, whereas the strength measure was designed to capture the amount of coupling between the joints (Figure 3). The fact that both measures were significant indicates that synergies affect both the strength of the coupling and its degree of intrusion during the movement. The increased inter-participant variability in the proportion measure suggests that this measure is more sensitive to individual coordination style compared to the strength measure.

Given the clear indications of intrusion of flexor synergy in the flexion coordination analyses (Figure 3), we would have expected to find a more robust task-dependent difference at the kinematics level (Table 3). This apparent discrepancy between the analyses may be driven by the fact that within synergy movements are sensitive to other deficits, such as weakness and loss of dexterity. The reduction in flexion-flexion coordination in the cup-to-mouth task, may be a marker of these additional contributors to the deficit.

Our results seemingly contradict those of Wagner and colleagues,^19^ by showing a contribution of flexor synergy intrusion to reaching abnormalities in the sub-acute stage. Wagner and colleagues compared the ability to make isolated joint movements in the acute post-stroke stage (all participants were tested within two weeks of their stroke) to performance in a reaching task. Their results showed that although the ability to individuate joints was impaired in the early sub-acute phase, it didn’t explain more than marginal portions of the variance of the reaching deficit (∼3%), while weakness explained the majority of variance (∼5-30%).^19^ In contrast, our study demonstrates that weakness and synergy together explain ∼53-71% of the variance of the FMA, ARAT, reaching extent, and peak velocity (Table 4), where each one of them explained at least 16% of the variance in kinematic performance measures (R > |0.41|) (Table 5). We argue that the discrepancy between these two studies relates to the kinematic measures used. The individuation index may relate more to the negative symptom of reduced dexterity, whereas our flexion-flexion measure may better capture the positive phenomenon of af flexor synergy. Thus, our measures may allow better disambiguation of synergy intrusion from more general joint coordination deficits.

Studies in the second half of the 20^th^ century that attempted to formally characterize changes in the post-stroke arm paretic phenotype over the course of recovery noted that in addition to weakness, stroke participants also suffer from obligatory flexor and extensor synergies.^5,6^ The Fugl-Meyer Assessment (FMA)^9^ was designed, in part, to capture such synergy intrusion. The FMA correlates with other measures of synergies^10,11,31^ and is abnormal in acute, sub-acute, and chronic stroke.^32–36^ Our measures also correlated with the FMA (Table 5). Surprisingly, despite the synergy-measurement rationale for the FMA, it can also be scored low because of weakness alone, as demonstrated by those participants that did not display intrusion of synergies but had low FMA scores (Figure 5). Indeed, various studies have shown that the FMA correlates with weakness.^7,31,37,38^

These results, along with previous ones, have several implications. First, they emphasize the importance of going beyond clinical scales toward the characterization of neurological deficits with fine-grained kinematic analyses of natural movements.^39^ Second, they suggest that the meaning of an abnormal FMA change between participants: in some, it might primarily reflect weakness, and in others, the intrusion of synergies. Thus, response to an intervention when measured by the FMA may mean very different things, for example, when administered in sub-acute versus chronic stroke.^40–42^ Third, the fact that weakness plays such an important role in the motor control deficits in the acute and sub-acute stages post-stroke may be the reason that the RST, which is critical to the generation of large forces,^43^ gets upregulated in the chronic phase.^44–46^ Essentially, a weakness problem, especially when severe, gets replaced by a synergy problem. Novel interventions, likely best instigated at the acute and subacute stages, are going to be needed to mitigate this zero-sum game trade-off.

### Limitations

One potential limitation of this study is that our cup-to-mouth task didn’t require as much shoulder flexion as the reaching task (see supplementary material). This could be why we couldn’t find consistent differences in how stroke participants performed both tasks. We think that this is unlikely due to the qualitative difference between the tasks (the reaching task requires primarily moving outside the flexor synergy: shoulder flexion and elbow extension, and the cup-to-mouth task does not). Furthermore, this limitation does not affect the flexor synergy strength measure since that was calculated in a continuous segment in which the shoulder was flexing.

We did not perform EMG, albeit purposefully. As stated in the introduction, while abnormal EMG co-activation may be considered a necessary condition for the emergence of synergies, isolating pathological synergies in non-isometric conditions, i.e., during voluntary movements, using EMG is challenging and has not been attempted to our knowledge. In addition, abnormal EMG does not inevitably result in abnormal kinematics.^14^ Our interest was more about whether the joint flexor synergy had a deleterious effect on reaching and less about its origin. Indeed, we would be hard-pressed to devise an alternative explanation for an unwanted flexor synergy that does not involve muscle co-activation.

## Data availability

Raw data were collected at Ben-Gurion University and Adi Negev Nahalat Eran in Israel. Derived data supporting the findings of this study are available from the corresponding author on request.

## Supporting information

Supp Mov. A and B

Supp Mov. A and B

## Acknowledgments

We thank Sandra Deluzio for the insightful clinical interpretation of the data.

We thank the recovering participants and families who agreed to participate in this study and acknowledge the medical and research staff members of Adi Negev Nahalat Eran who assisted with the entire operation.

---

**Supplementary figure 1.**
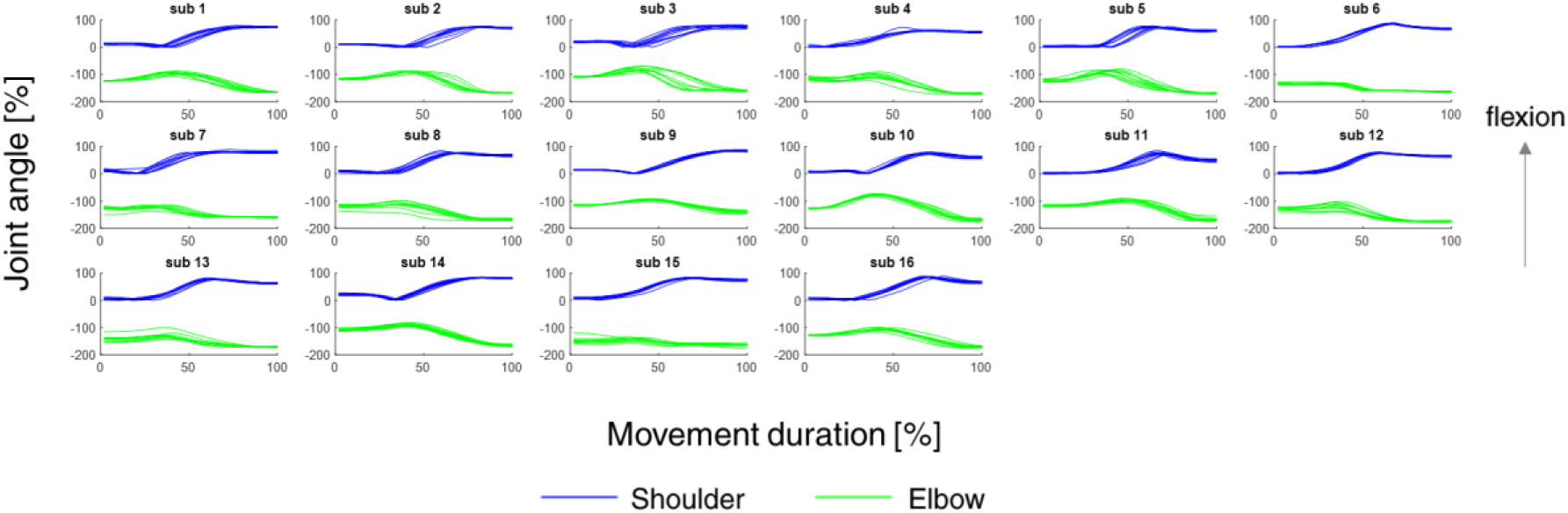
Shoulder and elbow angles of the non-dominant hand of controls in the reaching task. Angular trajectories of shoulder flexion (blue) and elbow flexion (green) of all control participants. Each movement iteration is represented as a single line of both shoulder and elbow trajectories.

**Supplementary figure 2.**
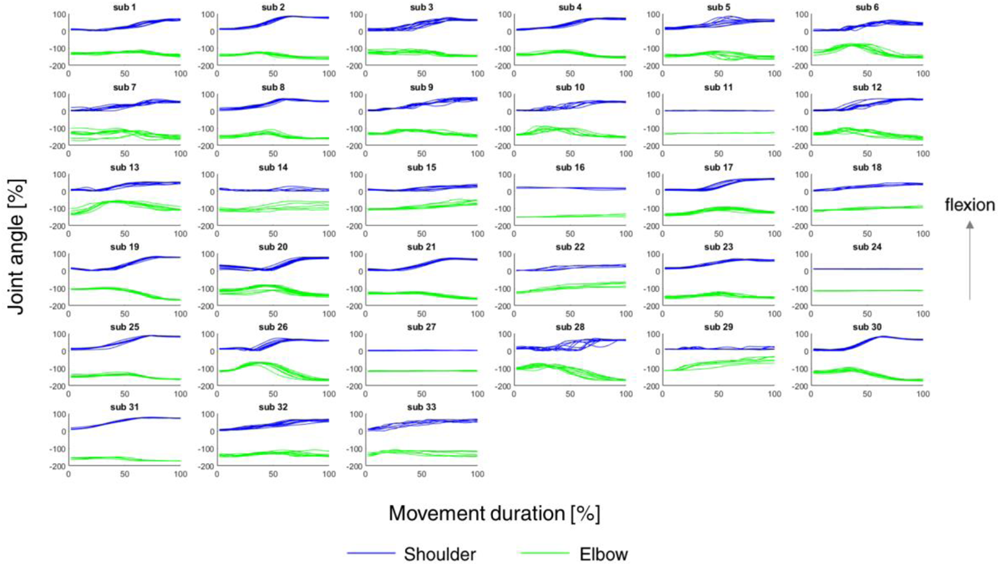
Shoulder and elbow angles of the paretic hand of sub-acute participants in the reaching task. Angular trajectories of shoulder flexion (blue) and elbow flexion (green) of all stroke participants. Each movement iteration is represented as a single line of both shoulder and elbow trajectories.

**Supplementary figure 3.**
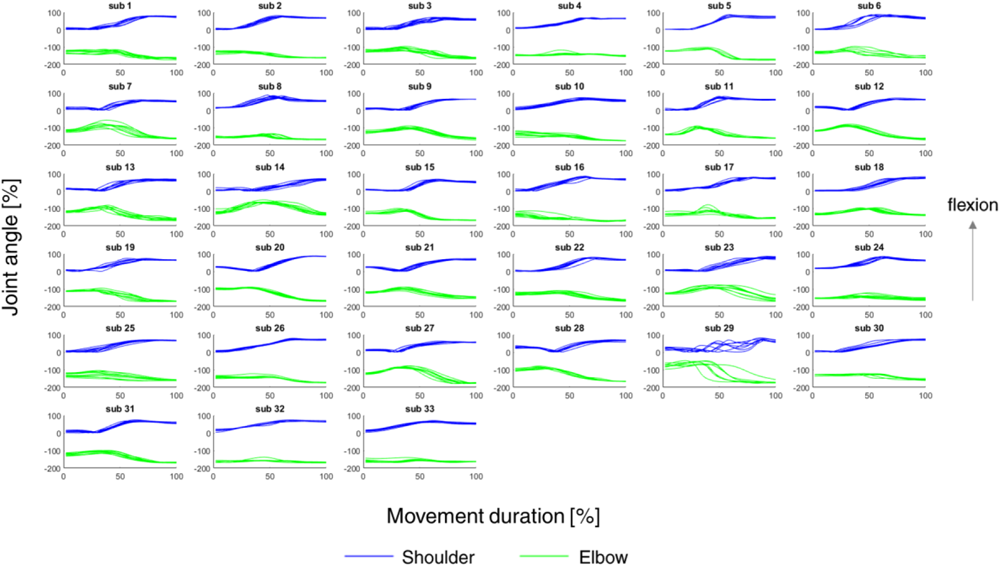
Shoulder and elbow angles of the non-paretic hand of sub-acute participants in the reaching task. Angular trajectories of shoulder flexion (blue) and elbow flexion (green) of all stroke participants. Each movement iteration is represented as a single line of both shoulder and elbow trajectories.

**Supplementary figure 4.**
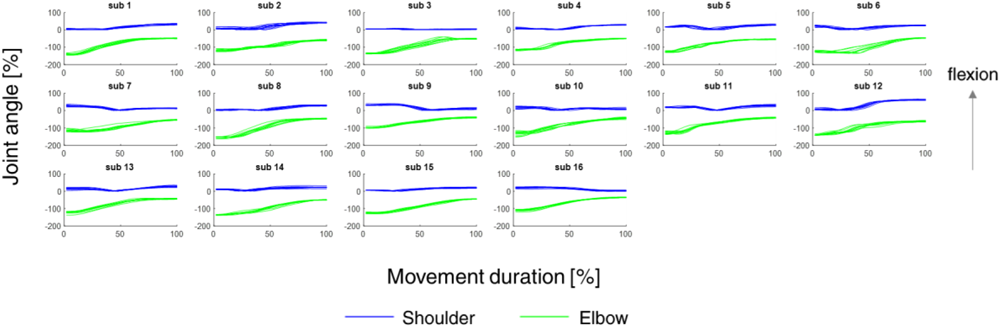
Shoulder and elbow angles of the non-dominant hand of controls in the cup-to-mouth task. Angular trajectories of shoulder flexion (blue) and elbow flexion (green) of all control participants. Each movement iteration is represented as a single line of both shoulder and elbow trajectories.

**Supplementary figure 5.**
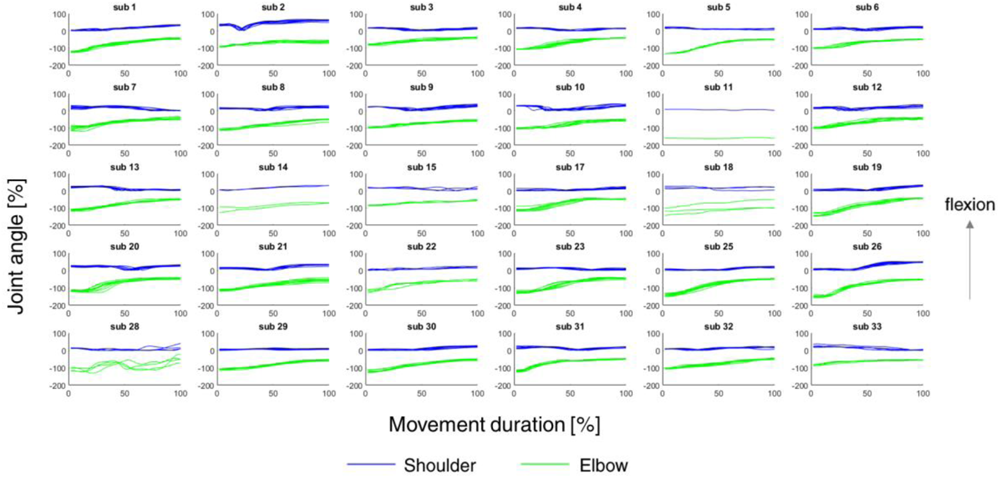
Shoulder and elbow angles of the paretic hand of sub-acute participants in the cup-to-mouth task. Angular trajectories of shoulder flexion (blue) and elbow flexion (green) of all stroke participants. Each movement iteration is represented as a single line of both shoulder and elbow trajectories.

**Supplementary figure 6.**
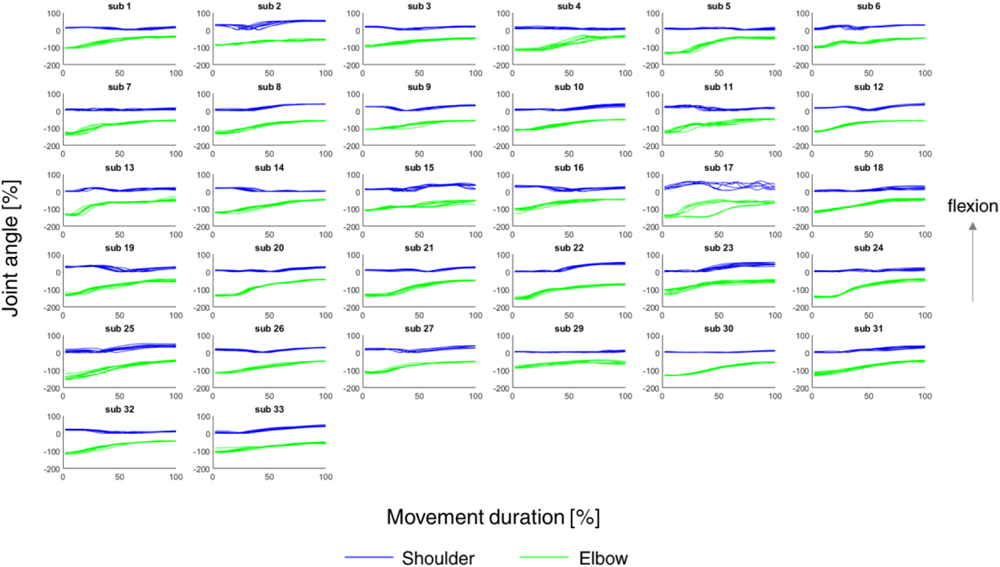
Shoulder and elbow angles of the non-paretic hand of sub-acute participants in the cup-to-mouth task. Angular trajectories of shoulder flexion (blue) and elbow flexion (green) of all stroke participants. Each movement iteration is represented as a single line of both shoulder and elbow trajectories.

